# Study Design and Interim Analysis of the Cancer Lifetime Assessment Screening Study in Canines (CLASSiC): The First Prospective Cancer Screening Study in Dogs Using Next-Generation Sequencing-Based Liquid Biopsy

**DOI:** 10.1101/2024.04.01.587600

**Authors:** Andi Flory, Suzanne Gray, Lisa M. McLennan, Jill M. Rafalko, Maggie A. Marshall, Kate Wotrang, Marissa Kroll, Brian K. Flesner, Allison L. O’Kell, Todd A. Cohen, Carlos A. Ruiz-Perez, Emily Sandford, Ana Clavere-Graciette, Ashley Phelps-Dunn, Rita Motalli-Pepio, Prachi Nakashe, Mary Ann Cristobal, Phadre Anderson, Susan C. Hicks, John A. Tynan, Kristina M. Kruglyak, Dana W. Y. Tsui, Daniel S. Grosu

## Abstract

**Objective:** The Cancer Lifetime Assessment Screening Study in Canines (CLASSiC) is a prospective, longitudinal cancer screening study, in which enrolled dogs are screened for cancer with physical exams and next-generation sequencing-based liquid biopsy testing on a serial basis. The goals of the first interim analysis, presented here, are to assess the benefits of using the OncoK9® liquid biopsy test as a cancer screening tool in a prospective clinical setting, and to demonstrate test performance for cancer detection, including preclinical detection.

**Subjects:** 726 presumably cancer-free client-owned dogs were prospectively enrolled in the study across 24 clinical sites in the US and Canada. Most subjects were at high risk of cancer at the time of enrollment based on age and/or breed. 419 dogs that were enrolled for at least one year and had at least two cancer screening study visits, or that had received a definitive or presumptive diagnosis of cancer up to the time of the interim analysis, were included in the analysis.

**Methods:** Clinical data and a blood sample were collected at each study visit (once or twice per year and when cancer was clinically suspected). Cell-free DNA extracted from plasma was tested by OncoK9® using next-generation sequencing (NGS) technology.

**Results:** 417 dogs were eligible for inclusion in the interim analysis and had classifiable outcomes, with a mean on-study duration of 422 days. Of these, 51 dogs were newly diagnosed with cancer (37 definitive, 14 presumptive), translating to a 12% (51/417) observed incidence over the study period; the liver, skin, bone, heart, spleen, lung, and lymph node(s) were the most common anatomic locations for disease. The prospectively observed sensitivity (detection rate) of the test was 56.9% (95% CI: 42.3-70.4%) with a specificity of 98.9% (95% CI: 97.0-99.6%). The prospectively observed positive predictive value was 87.9% (95% CI: 70.9-96.0%) and the negative predictive value was 94.3% (95% CI: 91.3-96.3%). NGS-based liquid biopsy doubled the overall number of cancer cases detected in this study population (from 25 to 51); remarkably, the detection rate for preclinical cancer was increased 4.6-fold from 12% (6/51) by routine care alone to 55% (28/51) by combining routine care with OncoK9® testing.

**Clinical Relevance:** CLASSiC is the first study to prospectively document the incidence of cancer in a predominantly high-risk canine population, and to prospectively demonstrate that the addition of NGS-based cancer screening to regularly scheduled wellness visits has the potential to substantially increase preclinical cancer detection in this population.

## INTRODUCTION

Cancer screening involves measures taken to find cancer in a patient with no clinical signs of the disease, at a time when the disease may be easier to treat.^1^ Historically, cancer screening in dogs has relied primarily upon the annual or semi-annual wellness visit; however, in most dogs cancer is detected after the development of clinical signs, often when the disease is advanced.^2^ Early detection through screening offers the best chance of improving outcomes for dogs with cancer.^3,4^

In 2021, a blood-based cancer detection test, OncoK9®, became clinically available for dogs. The test uses advanced sequencing technology (next-generation sequencing or “NGS”) to analyze free-floating fragments of DNA (cell-free DNA or “cfDNA”) for acquired (or “somatic”) genomic alterations that are associated with the presence of cancer.^5^ Referred to as “NGS-based liquid biopsy”, this test was clinically validated to detect 30 different cancer types, with a low false positive rate, in the CANcer Detection in Dogs (CANDiD) study.^6^ Since 2021, the test has been used clinically in thousands of dogs^7^, with the majority of samples submitted for screening of high-risk dogs with no current suspicion of cancer.^8^ A separate study, involving a subset of subjects from the CANDiD study, showed that NGS-based liquid biopsy has the ability to detect cancer preclinically, as well as at an early disease stage, making it a promising tool to increase early cancer detection when added to a wellness visit.^2^

NGS-based multi-cancer early detection (MCED) tests developed for use in humans over the past few years^9–11^ have benefited from large-scale observational studies demonstrating the ability of such testing to prospectively detect cancer in patients who were cancer-free at enrollment but were at high risk for the disease due to age (screening scenario),^12–14^ and in patients who presented with clinical signs suggestive of cancer (aid-in-diagnosis scenario).^15^ Such studies play an important role in documenting the real-world clinical utilization and performance of new tests.

The real-world performance of the OncoK9® test was previously studied in a retrospective review of 1,500 cases submitted for commercial testing, where sensitivity and specificity were found to be very similar to the CANDiD study.^8^ The current study, the Cancer Lifetime Assessment Screening Study in Canines (“CLASSiC”), is the first large-scale, multi-center, longitudinal cancer screening study in dogs executed prospectively according to a single protocol, with a rigorous outcomes data collection methodology including financial support to the sites for thorough case workups and documentation. In this study, serial whole blood samples and contemporaneous clinical data are collected at regular intervals from client-owned dogs that are presumably cancer-free at the time of enrollment, in order to detect cancer prior to the onset of clinical signs using a validated NGS-based liquid biopsy test (OncoK9®; PetDx, La Jolla, CA). Results are returned to the study sites in real time, with collection of outcome data on all cases. The study’s primary goal is to assess the benefits of using this test as a cancer screening tool in a prospective setting, with documentation of test performance in a first interim analysis. Long-term goals are: to determine the optimal interval for canine cancer screening; to evaluate the additional testing required to confirm a diagnosis following a positive OncoK9® result; to evaluate owner satisfaction and attitudes toward future screening visits; and, to evaluate veterinarian satisfaction with this screening test and its impact on patient management.

Presented here are the design of the CLASSiC study as well as selected results from the first interim analysis of the study.

## METHODS

### Cohorts

Enrollment for CLASSiC started in December 2021 under two IACUC-approved protocols: CLASSiC and CLASSiC Plus. CLASSiC patients were enrolled at primary care practices, and CLASSiC Plus patients were enrolled at primary care sites that were partnered with a participating specialist hospital site, or were enrolled at an academic institution. Enrollees in CLASSiC and CLASSiC Plus originated from two cohorts. Cohort 1 consisted of dogs previously enrolled as “presumably cancer-free” in the control arm of the CANDiD study^6^ and were determined to still be presumably cancer-free at the time of enrollment in CLASSiC/CLASSiC Plus. Cohort 2 consisted of newly enrolled dogs that were age 7 and older or that belonged to breeds at higher lifetime risk of cancer,^16,17^ who could enroll at age 4 and older.

### Inclusion and Exclusion Criteria

The following inclusion criteria were required for enrollment of dogs in Cohorts 1 and 2: must be presumably cancer-free (i.e., not known to have cancer, not currently suspected to have cancer, and no prior history of resolved cancer); safe for 14-17 mL blood draw without fluid replacement; owner willing and able to provide written informed consent; owner agreed to comply with all study procedures and to provide all related study information; and owner agreed to make their best effort to notify the study site if their dog developed cancer at any time during or after study participation (in the event of early withdrawal).

Exclusion criteria were as follows: dogs with a previous diagnosis of cancer or currently suspected/known to have cancer; dogs that had experienced physical trauma (including surgery or biopsy) within 7 days prior to blood collection; dogs that were pregnant; dogs with severe anemia (defined as hematocrit <18% with body weight <7.3 lb/3.3 kg); dogs that had received a donor-derived bone marrow transplant or donor-derived stem cell therapy at any time; dogs that had received an organ transplant; dogs that had received a whole blood transfusion within the past 3 months; non-domestic canids, hybrids, chimeras, or mosaics; and, dogs for which collection of whole blood provided an unacceptable risk to the site staff and/or the dog. Dogs with suspected common benign cutaneous/subcutaneous tumors (such as lipomas and skin tags) were not excluded. Dogs with acute or chronic medical conditions/diagnoses were eligible as long as cancer was not suspected. While physical trauma including surgery or biopsy within the previous 7 days were exclusionary, dogs that had routine blood draws, fine needle aspiration (FNAs), or cystocentesis within 7 days prior to blood collection were eligible, as these were not considered to cause significant trauma.

### Enrollment Visit

At the enrollment visit, informed consent and medical history (including vaccination history for the past 3 months) were collected from the owner, and the following information was documented: breed(s) as reported by owner, sex, spay/neuter status (including age at spay/neuter, if known), and age (or approximation). The owner also completed a “Common Cancer Signs Questionnaire” and any abnormalities marked by the owner were assessed as “Clinically Significant” or “Not Clinically Significant” by the site’s principal investigator. A complete physical exam, including oral and rectal exams and lymph node and neck palpation, was conducted, and abnormal findings were noted, by the managing veterinarian.

Following the exam, whole blood (14.0 – 17.0 mL per subject) was collected using Cell-Free DNA Collection tubes (Roche Diagnostics; Rotkreuz, Switzerland), with no in-clinic processing (e.g., refrigeration, freezing, centrifugation or separation) required. Samples were shipped to the central testing laboratory (PetDx; La Jolla, CA) at ambient temperature.

### Routine Follow-Up Visits

Dogs in Cohort 1 had Routine Follow-Up Visits (RFUVs) at a minimum cadence of once per year (+/-30 days), with the option of semi-annual testing (i.e., every 6 months) at the owner’s discretion. Dogs in Cohort 2 had semi-annual RFUVs (i.e., every 6 months, +/-30 days). At each RFUV, a complete physical exam (including oral/rectal exam and lymph node/neck palpation) was conducted, the dog’s weight was recorded, medical and vaccine history was updated, the owner completed the Common Cancer Signs Questionnaire, and whole blood was collected (as described above).

### Suspicion of Cancer Visits

In addition to the scheduled Enrollment visits and RFUVs, managing veterinarians were permitted to submit blood samples from unscheduled visits triggered by a suspicion of cancer (Suspicion of Cancer Visits, SOCVs), based on clinical presentation or results of laboratory tests or imaging performed outside of regularly-scheduled study visits. In the case of an SOCV, a *Cancer Signal Detected (CSD)* result prompted a confirmatory cancer evaluation as described below.

### Sample Testing and Results Delivery

All blood samples were sent to a central laboratory for processing and underwent NGS analysis using the OncoK9® test, as previously described.^6,8^ In the case of a *Cancer Signal Not Detected* (*CSND*) result, the dog returned in 6 or 12 months for their next scheduled RFUV. If an enrollee returned to the principal investigator (PI) for a SOCV, a blood sample was obtained and visit details (exam and diagnostic findings) were documented. In the case of an *Indeterminate* or *Sample Failure* OncoK9® result, the dog returned for a Retest Visit within 4 weeks of the last test date.

### Follow-Up Procedures

In the case of a *CSD* result, the dog returned for a Confirmatory Follow-Up Visit (CFUV). At this visit, the owner completed a Common Cancer Signs Questionnaire, the medical/vaccine history was updated, a thorough physical exam was performed, and weight was documented. Additionally, a confirmatory cancer evaluation (CCE) to evaluate the most common cancer locations in the body, was recommended as part of a comprehensive clinical work-up. The cost of the CCE was covered by the study Sponsor (PetDx; La Jolla, CA), and could include: CBC; serum chemistries and urinalysis; 3-view thoracic radiographs and radiographs of any areas of focal pain or lameness, with written review by a board-certified radiologist; abdominal ultrasound performed by a board-certified radiologist or internist with a complete written report; fine needle aspiration and cytology of any palpable mass(es) measuring >1 cm and/or enlarged lymph nodes, submitted to the site’s veterinary diagnostic laboratory.

The PI and/or managing veterinarian determined the appropriate diagnostic workup, and the study did not require any procedures outside of each site’s standard of care. All exam and diagnostic reports were provided to the Sponsor. Additional consultation regarding how to proceed based on the result of the CCE and Confirmatory Follow-Up OncoK9® test, was provided by clinical study support veterinarians (board-certified in Internal Medicine or Oncology) employed by the Sponsor.

### Additional Follow-Up for CLASSiC Plus Subjects

A clinical cancer diagnosis following a *CSD* result and a CCE in a CLASSiC Plus enrollee resulted in further assessment for staging (e.g., additional imaging, echocardiogram, molecular diagnostics, bone marrow/organ cytology, and/or special staining), all of which were covered by the Sponsor.

The owner, in consultation with the clinician, could then decide to pursue surgery and/or treatment. Surgery and/or treatment was not dictated by the Sponsor and, if elected, was the financial responsibility of the owner. Upon diagnostic resolution, the owner received an OncoK9® Satisfaction Questionnaire via email from the Sponsor.

### Owner and Veterinarian Satisfaction Surveys

Following receipt of liquid biopsy results from the enrollment visits and RFUVs, and following confirmation of a cancer diagnosis in their dog (if applicable), an OncoK9® Owner Satisfaction Questionnaire was sent via email by the Sponsor. Completion of this electronic survey was voluntary and did not impact eligibility for participation in the study.

Additionally, OncoK9® Veterinarian Satisfaction Questionnaires were periodically sent by the Sponsor via email to the managing veterinarians. Completion of this electronic survey was voluntary and did not impact eligibility for participation in the study.

### Interim Analysis

The following criteria were required for inclusion in this first interim analysis of the CLASSiC study: 1) the dog had been enrolled for at least 1 year and had completed 2 or more assessments with liquid biopsy testing (e.g., enrollment visit plus at least one RFUV); or 2) the dog had been enrolled in CLASSiC for less than 1 year, but had received a definitive or presumptive diagnosis of cancer while on the study.

One of three “diagnostic tiers” was assigned to each dog with cancer based on how the diagnosis was achieved. Tier 1 indicated a definitive diagnosis microscopically confirmed by pathologist interpretation of a histopathology or cytology sample. Tier 2 indicated a presumptive diagnosis not microscopically confirmed but strongly suspected due to direct visualization or imaging (interpreted by a radiologist or board-certified specialist), strongly suspected on histopathology or cytology but not definitively diagnosed, or made on cytology but not confirmed by a pathologist (i.e., in-house cytology). Tier 3 indicated a presumptive diagnosis based on strong clinical suspicion alone. Herein, Tier 1 diagnoses are referred to as “definitive” and Tier 2 and 3 diagnoses are referred to as “presumptive”.

Test sensitivity is defined as the percentage of all cancer-diagnosed dogs who received a *CSD* (positive) result by OncoK9®. Test specificity is defined as the percentage of all presumably cancer-free dogs who received a *CSND* (negative) result by OncoK9®. Observed positive predictive value (PPV) refers to the percentage of dogs who received a *CSD* result by OncoK9® and were subsequently diagnosed with cancer. Observed negative predictive value (NPV) refers to the percentage of dogs who received a *CSND* result by OncoK9® and were presumably cancer-free at the conclusion of the interim analysis.

All metrics are summarized as median and range, unless otherwise stated. All confidence intervals reported are two-sided 95% binomial intervals.

## RESULTS

### CLASSiC Enrollee Demographics

As of the time of database lock for this first interim analysis (December 31, 2023), 726 dogs had been enrolled in CLASSiC/CLASSiC Plus across 24 sites in the US and Canada (21 general practices, 2 academic sites, and 1 specialty center). Subjects ranged in age from 1 to 20 years (median 10.5 years); weights ranged from 2.1 to 87.1 kg (median 44.6 kg); 50% (n=360) of subjects were male and 50% (n=366) were female; 96% of all subjects spayed or neutered; 61% (n=446) of subjects were purebred and 39% (n=279) mixed-breed, with one additional subject with missing breed classification.

### Interim Analysis Subjects and Samples

Of the 726 enrollees, 58% of subjects (n= 419) met inclusion criteria for the interim analysis, having been in the study for a mean time of 422 days (median = 385 days). The remaining subjects did not meet interim analysis inclusion criteria or were removed from analysis for one or more of the following reasons: study site closed; lost to follow-up; owner withdrew consent; owner moved; protocol violation; and death, unknown cause.

At the time of the interim analysis, a retrospective review of subject demographics was performed in order to determine individual cancer risk at the time of enrollment, based on age, breed, and/or weight. Dogs were considered at *increased risk for cancer* based on criteria defined by Rafalko et al^18^: any dog 7 years of age or older, dogs at or above the recommended age to start cancer screening based on breed, and dogs at or above the recommended age to start cancer screening based on weight for mixed breeds or uncommon breeds. Based on this review, eighty-four percent (350/419) of subjects were categorized as being at increased risk for cancer.

At interim analysis, a total of 1,871 samples had been submitted for OncoK9® testing from these 419 subjects. Only *CSD* and *CSND* results were used to calculate test performance. A binary *CSD* or *CSND* result was issued for 96% of samples (n=1,797); 2.5% (n=46) of samples were *Sample Failures* (19 due to insufficient blood volume, 13 due to transit issues, 9 due to sequencing-related or laboratory processing issues, and 5 due to expired tubes) and required repeat testing by OncoK9®; 1.5% (n=28) of results were *Indeterminate* and required repeat testing by OncoK9®.

### Outcome Data and Test Performance

Of the 419 subjects in the interim analysis, 417 had classifiable outcomes (i.e., had sufficient clinical data to be assigned a final outcome classification of true positive, false positive, true negative, or false negative); the remaining 2 subjects had *CSD* results with confirmatory cancer evaluations in progress at the time of database lock.

Of the subjects with classifiable outcomes, 51 were newly diagnosed with cancer (37 definitive, 14 presumptive) during the study period, translating to a 12% (51/417) incidence of disease in this population. Ninety-four percent (48/51) of the cancer-diagnosed dogs were at increased risk for cancer (based on the above criteria) at the time of diagnosis. Cancer types/locations identified in these patients are summarized in **Table 1**, with the liver, skin, bone, heart, spleen, lung, and lymph node(s) as the most common anatomic locations.

**Table 1:** Cancer types/locations for the 51 cancer-diagnosed dogs in the first interim analysis of the CLASSiC study

| Definitive Diagnoses (n=37) |  |  |  |  |
| --- | --- | --- | --- | --- |
| Histologic type | Anatomic location | # of cases | #TP | #FN |
| Hemangiosarcoma | Heart | 1 | 1* | 0 |
|  | Liver + spleen | 2 | 2* | 0 |
|  | Spleen | 1 | 1* | 0 |
| Hepatocellular carcinoma | Liver | 3 | 2* | 1 |
| Histiocytic sarcoma | Joint | 1 | 1 | 0 |
|  | Liver + lung | 1 | 1* | 0 |
| Leiomyosarcoma | Liver | 1 | 1* | 0 |
| Leukemia, Chronic Lymphoid | Blood | 1 | 0 | 1 |
| Lymphoma, epitheliotropic | Skin | 1 | 0 | 1 |
| Lymphoma, intermediate to large cell | Lymph node(s) | 2 | 2* | 0 |
| Lymphoma, intermediate to large cell + suspect hemangiosarcoma | Lymph node(s) + Spleen/Liver/Lung | 1 | 1* | 0 |
| Mammary ductular carcinoma | Mammary | 1 | 0 | 1 |
| Mast cell tumor | Skin | 5 | 0 | 5 |
| Mast cell tumor + suspect hemangiosarcoma | Skin + Heart | 1 | 1* | 0 |
| Meningioma | Brain | 1 | 1 | 0 |
| Osteosarcoma, appendicular | Scapula | 1 | 1* | 0 |
|  | Humerus | 1 | 1 | 0 |
|  | Femur | 1 | 0 | 1 |
|  | Tibia | 1 | 0 | 1 |
| Pheochromocytoma | Adrenal gland | 1 | 1* | 0 |
| Plasmacytoma | Skin | 1 | 0 | 1 |
| Prostatic carcinoma | Prostate | 1 | 1* | 0 |
| Sarcoma | Mandible | 1 | 1* | 0 |
|  | Digit | 1 | 0 | 1 |
|  | Cecum | 1 | 0 | 1 |
| Sertoli cell tumor | Testicle | 1 | 0 | 1 |
| Soft tissue sarcoma | Skin | 1 | 0 | 1 |
|  | Retrobulbar | 1 | 0 | 1 |
| Undefined subtype | Muscle | 1 | 1* | 0 |
| **Presumptive Diagnoses (n=14)** |  |  |  |  |
| **Histologic type** | **Anatomic location** | **# of cases** | **#TP** | **#FN** |
| NA | Brain | 1 | 0 | 1 |
| NA | Heart (suspect hemangiosarcoma) | 3 | 2* | 1 |
| NA | Heart/liver/lung/spleen (suspect hemangiosarcoma) + liver (suspect hepatocellular carcinoma) | 1 | 1* | 0 |
| NA | Heart/spleen (suspect hemangiosarcoma) | 1 | 0 | 1 |
| NA | Liver (suspect hepatocellular carcinoma) | 2 | 2* | 0 |
| NA | Liver/spleen (suspect hemangiosarcoma) | 1 | 1* | 0 |
| NA | Lung (suspect histiocytic sarcoma) | 1 | 1* | 0 |
| NA | Lymph node(s) (presumptive lymphoma) | 1 | 1* | 0 |
| NA | Polyostotic (suspect multiple myeloma) | 1 | 0 | 1 |
| NA | Prostate | 1 | 1* | 0 |
| NA | Skin (presumptive mast cell tumor) | 1 | 0 | 1 |

\*Detected by OncoK9® testing in at least one dog at a scheduled study visit (enrollment visit or routine follow-up visit). TP = True Positive. FN = False Negative. NA = Not Available.

*Cancer Signal Detected* (positive) OncoK9® results were issued for 29 of these 51 dogs, for a sensitivity (detection rate) of 56.9% (95% CI: 42.3-70.4%). There were 366 dogs with no cancer diagnosed during the study period. *Cancer Signal Not Detected* (negative) OncoK9® results were issued for 362 of these dogs, for a specificity of 98.9% (95% CI: 97.0-99.6%). (**Table 2**)

**Table 2:** Clinical performance characteristics of the OncoK9® test in subjects with classifiable outcomes (n=417) in the first interim analysis of the CLASSiC study

|  |  | Cancer status |  |  |
| --- | --- | --- | --- | --- |
|  |  | Present<br>(n=51) | Absent<br>(n=366) |  |
| OncoK9®<br>result | *Cancer Signal Detected*<br>(n=33) | True Positives<br>29 | False Positives<br>4 | Observed PPV<br>87.9%<br>(95% CI: 70.9-96.0%) |
|  | *Cancer Signal Not Detected*<br>(n=384) | False Negatives<br>22 | True Negatives*<br>362 | Observed NPV<br>94.3%<br>(95% CI: 91.3-96.3%) |
|  |  | Observed Sensitivity<br>56.9%<br>(95% CI: 42.3-70.4%) | Observed Specificity<br>98.9%<br>(95% CI: 97.0-99.6%) |  |

\*Assumed true negatives based on available information supporting the absence of cancer at the time of outcome data collection.

In total, 33 dogs received *Cancer Signal Detected* results, of which 29 were diagnosed with cancer, corresponding to an observed PPV of 87.9% (95% CI: 70.9-96.0%); 384 dogs received *Cancer Signal Not Detected* results, of which 362 had no diagnosis of cancer during the study period, corresponding to an observed NPV of 94.3% (95% CI: 91.3-96.3%). (**Table 2**)

### True Positives

There were 29 dogs with true positive results from liquid biopsy testing (i.e., had a *CSD* OncoK9® result and were diagnosed with cancer); of these, 20 diagnoses were definitive and 9 were presumptive (representing 7 anatomic locations; **Table 1**); furthermore, 34% (n=10) had their first *CSD* result at their enrollment visit, 52% (n=15) had their first *CSD* result at a RFUV, and 14% (n=4) had their first *CSD* result at a SOCV. In total, 86% of true positive dogs (25/29) had their first *CSD* result at a regularly-scheduled study visit (i.e., enrollment visit or planned RFUV). In these dogs, the time to diagnostic resolution (i.e., the time elapsed from the date of the first *CSD* result reported to the managing veterinarian to the date of cancer diagnosis) ranged from 4 – 259 days, with a median of 23 days; 56% of these dogs (14/25) achieved diagnostic resolution within one month, 80% (20/25) within 3 months, and 92% (23/25) within 6 months. In the remaining two patients (one with a meningioma and one with osteosarcoma of the scapula), diagnostic resolution was achieved after 188 and 259 days (respectively).

A review of records for the 29 dogs with true positive results showed that cancer was first localized by physical exam in 11 dogs, abdominal ultrasound (AUS) in 10 dogs, thoracic radiographs (CXR) in 3 dogs, a combination of AUS and CXR in 1 dog, limb radiographs in 1 dog, and advanced imaging in 2 dogs (echocardiogram and MRI, respectively). A definitive diagnosis was reached in 20 of these 29 dogs; 12 diagnoses were made via histopathology and 8 via cytology. In the cases confirmed via histopathology, tissue samples were obtained via the following methods: 4 necropsy, 3 splenectomy (one case also included liver biopsy, and one other included liver lobectomy), 1 liver lobectomy, 1 joint mass debulking excisional biopsy, 1 scapulectomy, 1 craniotomy, and 1 adrenalectomy + cavotomy. In the cases confirmed via cytology, 6 did not require ultrasound guidance, 1 involved US-guided FNA of the liver, and 1 involved US-guided FNA of the prostate.

### False Positives

There were 4 dogs with false positive results from liquid biopsy testing (i.e., had a *CSD* OncoK9® result and were not diagnosed with cancer prior to database lock for this interim analysis). All had routine labs (CBC, blood chemistry, urinalysis), CXR and AUS. Additional diagnostics included: echocardiogram (n=2), FNA of cutaneous/subcutaneous nodules (n=2), FNA of liver nodule (n=1), and thoracic CT scan (n=1). All 4 subjects remain under observation in the CLASSiC study.

### False Negatives

There were 22 dogs with false negative results from liquid biopsy testing (i.e., had a *CSND* OncoK9® result and were diagnosed with cancer prior to database lock for this interim analysis); 17 of these diagnoses were definitive, and 5 were presumptive (**Table 1**). The time from the patient’s last *CSND* liquid biopsy result to the time of cancer diagnosis ranged from 0 to 250 days, with 4 of these diagnoses occurring more than 6 months after the patient’s last *CSND* result (i.e., 192, 196, 235, and 250 days); additionally, 8 of these diagnoses involved small (<2 cm) cutaneous tumors.

### Cancer Detection in Relation to Clinical Signs

Review of records for the 51 cancer-diagnosed dogs found that only 12% (n=6) had cancer detected preclinically solely due to findings at a wellness visit, or during other routine care (not due to liquid biopsy testing); *ad hoc* visits for suspicion of cancer (SOCVs) detected cancer in another 37% (n=19) of these dogs, triggered solely by the manifestation of clinical signs (not due to liquid biopsy testing).

Cancer was detected preclinically due to a workup prompted solely by a *CSD* OncoK9® result in another 43% (n=22) of these dogs; and an additional 8% (n=4) had clinical signs suggestive of cancer, but were diagnosed only after a confirmatory cancer evaluation was prompted by a *CSD* result. In summary, 49% (n=25) of cancer-diagnosed dogs in this interim analysis had their cancer detected by various means that did not involve liquid biopsy testing, and 51% (n=26) had their cancer detected due to liquid biopsy testing.

### Safety

Of the 726 dogs enrolled in CLASSiC, 35 had reported adverse events as a direct result of the blood draw for OncoK9® testing: 32 dogs experienced a hematoma or bruising at the venipuncture site, 2 dogs were anxious during blood collection, and 1 dog required two venipunctures to obtain the sample. There were no adverse events reported due to diagnostic workups prompted by *CSD* liquid biopsy results.

### Owner Satisfaction

As of December 31, 2023, a total of 1,116 OncoK9® Satisfaction Questionnaires were sent to owners, and 343 responses were received – a response rate of 31%. On a scale from 0 (strongly disagree) to 10 (strongly agree), when asked their level of agreement with the statement “annual/semi-annual screening with the OncoK9® test is a good thing for my pet”, the average score from respondents was 9.3±1.5; for “detecting cancer early in my dog, when treatment options are better, is very important to me”, the average score was 9.6±1.1; and for “overall, I am satisfied with the OncoK9® screening test”, the average score was 9.3±1.6. When asked “how likely are you to pay for your pet’s annual screening with OncoK9® as recommended by your veterinarian” the following responses were noted: very unlikely (n=3), unlikely (n=43), likely (n=184), very likely (n=113). When asked “how likely are you to recommend cancer screening by OncoK9® to friends and family who own dogs”, the following responses were noted: very unlikely (n=2), unlikely (n=19), likely (n=179), very likely (n=143).

### Veterinarian Satisfaction

As of December 31, 2023, a total of 44 surveys were sent to managing veterinarians, and 10 responses were received – a response rate of 23%. On a scale from 0 to 10, when asked their level of agreement with the statement “OncoK9® reports are easy to navigate and understand”, the average score from respondents was 9.7±0.6; for “I feel prepared to present the OncoK9® results to my clients”, the average score was 9.6±0.8; for “OncoK9 helps with the clinical management of my study patients”, the average score was 9.5±0.8; for “overall, I am satisfied with the OncoK9 test as a multi-cancer screening tool in dogs who may be at higher risk for developing cancer due to their age and/or breed”, the average score was 8.8±1.8; and for “how likely are you to recommend OncoK9® to fellow colleagues/other veterinarians as a multi-cancer screening tool in dogs who may be at higher risk for developing cancer due to their age and/or breed”, the average score was 8.9±1.6.

## DISCUSSION

CLASSiC is the first large-scale, multi-center, longitudinal study to prospectively evaluate the utilization and performance of serial cancer screening with NGS-based liquid biopsy in dogs. In this first interim analysis, OncoK9® testing detected genomic alterations (a cancer signal) in 56.9% of dogs who had received a definitive or presumptive diagnosis of cancer during the study period. The observed specificity of the test was 98.9%, corresponding to a 1.1% false positive rate. In the cohort of dogs with true positive results, a variety of cancer diagnoses were represented, with most occurring internally (e.g., heart, lungs, liver, spleen) in locations that may be difficult to detect by physical exam alone, highlighting the utility of a blood-based cancer detection test. Importantly, following a *CSD* result in these patients, cancer was often localized or diagnosed within a matter of weeks (median, 23 days) using evaluation methods commonly available in a general practitioner’s office (e.g., comprehensive physical exam, CXR, and FNA cytology without ultrasound guidance). A recent real-world cohort tested with the same technology^8^ demonstrated an even shorter time to diagnostic resolution (median, 11 days); the discrepancy may be due to additional requirements in the present study, such as for board-certified radiologist ultrasound and imaging reports.

In human medicine, the National Cancer Institute collects incidence, prevalence, and survival data for cancer patients across the United States through the Surveillance, Epidemiology and End Results (SEER) registry; however, no such organization exists in veterinary medicine.^19^ Furthermore, registries that collect data on cancer incidence and mortality of dogs are not standardized, making comparisons among them challenging^20^; and they provide little publicly available data.^19^ As a result, the precise incidence of cancer in dogs remains to be established.^21^ CLASSiC is the first study to prospectively document the incidence of cancer in a predominantly high-risk canine population; the incidence over the period covered in this first interim analysis (mean = 422 days; median = 385 days) was observed to be 12%. This is consistent with, and possibly higher than, the 8-10% annual incidence of cancer in dogs previously estimated in the literature.^6^

The most common anatomic locations where cancer was found or was strongly suspected were: liver, skin, bone, heart, spleen, lung, and lymph node(s). The apparent increase in cancer incidence observed in this cohort may be a form of The Will Rogers phenomenon^22^, also known as “stage migration”. By using more sensitive diagnostic techniques, more cases are detected, and prognosis for stage-specific cohorts improves, given that many of the additional cases are detected at earlier stages. This phenomenon has been described in dogs with lymphoma, wherein the use of more sensitive staging tests detects more disease.^23^ A longer-term objective of CLASSiC is to determine whether the addition of NGS-based liquid biopsy to wellness visits improves stage-specific prognosis; this first interim analysis did not explore this hypothesis.

Although managing veterinarians were permitted to submit samples from unscheduled visits prompted by a clinical suspicion of cancer (SOCVs), the majority of dogs in this study with a true positive result (25/29, 86%) had their first *CSD* result at a regularly-scheduled study visit. Overall, this represents 49% (25/51) of all dogs diagnosed with cancer in this study. This is in contrast to the cancer detection paradigm in current clinical practice: a recent analysis of >350 dogs with cancer found that only 4% had their disease detected at a routine wellness visit.^2^

Of the 51 dogs diagnosed with cancer during the study period, 22 had false negative results from NGS-based liquid biopsy testing. As previously discussed, the detection rate of this type of testing varies by cancer type due to a variety of potential factors, including tumor size, extent of disease, and anatomic location.^6^ For example, aggressive cancers (such as lymphoma, hemangiosarcoma, and osteosarcoma) tend to be more readily detectable than anatomically isolated cancers (such as malignancies of the central nervous system, urinary bladder, or prostate).^6^ Additionally, NGS-based liquid biopsy is not indicated for the evaluation of cutaneous or subcutaneous masses that are typically detectable by physical examination and readily accessible for FNA or biopsy.^6^ Analysis of the 22 false negative cases in this interim analysis found that 36% (8/22) of these patients had small (<2 cm) cutaneous tumors.

When evaluating *why* cancer was ultimately detected in the 51 cancer-diagnosed dogs in this study, physical exams and other investigations (performed at a routine wellness visit, during other routine care, or at off-schedule sick visits) ultimately resulted in the diagnosis of 25 cases, whereas workups prompted by *CSD* results from NGS-based liquid biopsy ultimately resulted in the diagnosis of 26 additional cases of cancer. In summary, NGS-based liquid biopsy doubled the number of cancer cases detected in this study population.

Overall, 6 of the 51 cancer cases were detected preclinically solely due to findings during a wellness visit or other routine care, consistent with a previously reported observation.^2^ An additional 22 of the 51 cancer cases were detected preclinically due to a workup prompted solely by a *CSD* OncoK9® result.

Therefore, the preclinical cancer detection rate increased 4.6-fold from 12% (6/51) by routine care alone to 55% (28/51) by combining routine care with NGS-based liquid biopsy, confirming that adding blood-based cancer screening to regularly scheduled canine wellness visits has the potential to substantially increase preclinical cancer detection.

Overall, owner and veterinarian experience with this type of testing was very positive, although the veterinarian cohort had a much smaller number of returned questionnaires available for analysis.

The data reported in this study are the latest in a series of studies published by our group on NGS-based liquid biopsy for cancer detection in dogs.^6,8,24,25^ Non-significant differences in sensitivity (p=.460) and specificity (p=.338) were found across this interim analysis, the test’s clinical validation study^6^, and a third study involving real-world patients tested by the same method.^8^ The consistent performance observed across these three large independent studies underscores the robust design and stable technical characteristics of the test over time, as well as the value of having employed independent training and testing sets in the initial clinical validation^6^ of the OncoK9® test – an advanced study design strategy intended to reduce the risks of data overfitting and consequent overestimation of actual test performance.^9^

Early detection (whether in the preclinical period or at an earlier stage of disease) has been extensively documented to improve outcomes for many cancer types in humans^26^ as well as in dogs.^27–46^ A significant concern with all types of cancer screening is the potential for a large number of false positive results, which could lead to unnecessary procedures and expense as well as emotional distress.^5^ The high PPV prospectively observed in this study provides reassurance that most OncoK9® *CSD* results in a high-risk cancer screening population are in fact true positives, and cancer will be found if an adequate confirmatory evaluation is performed. At 87.9%, the PPV observed in this interim analysis is twice as high as the PPV of 43.1% recently documented in the landmark PATHFINDER study^13^ that enrolled over 6,000 people at high risk of cancer; while the validated performance of the two NGS-based MCED tests (OncoK9® and Galleri®) is very similar in terms of sensitivity and specificity,^6,8,9^ the annual incidence of cancer is much higher in dogs compared to humans,^6,8,47^ largely accounting for the higher PPV observed in the CLASSiC study.

The current study has certain limitations. The findings reported in this interim analysis represent only a subset of initial results from the CLASSiC study, gathered over a relatively short observation period; further study of this cohort is expected to provide more robust evidence regarding the performance and clinical utility of NGS-based cancer screening in dogs, including the cumulative detection rate afforded by repeat interval testing over many years of life.^6^ A longer period of observation is needed to determine the appropriate cadence of screening with liquid biopsy (i.e., annual vs. semi-annual) and to address the question of whether clinical outcomes in the high risk cancer screening population are improved as a result of earlier detection. Other limitations include the fact that the owner and veterinarian surveys did not use validated scales, and that the pet parents and veterinary clinics that enroll in a study such as this (and choose to respond to a voluntary survey) may represent a biased population. Finally, managing veterinarians and owners were not required to pursue confirmatory testing after a *CSD* result, so the opportunity to arrive at a presumptive or definitive cancer diagnosis was limited in some cases.

The ability to increase preclinical cancer detection at a dog’s wellness visit is a goal shared by clinicians and pet owners alike. Multi-cancer early detection via NGS-based liquid biopsy offers great potential to achieve this goal, using a simple blood collection that can be performed in any clinical setting. CLASSiC, the first lifetime cancer screening study in dogs, is positioned to provide the necessary evidence to support changes in clinical practice and guidelines-based recommendations, and represents a significant milestone in veterinary medicine.

## Acknowledgements

The authors thank all veterinarians and veterinary caregivers that contributed to this study, who talked to their clients about the importance of early cancer detection, and who see the need for a proactive, not reactive, approach to cancer detection. The authors also thank all the pets and pet parents who participated in this groundbreaking study, for their incredible dedication to science.

